# A framework for analysis of real-time nucleic acid amplification data using novel multidimensional standard curves

**DOI:** 10.1101/379180

**Authors:** Ahmad Moniri, Jesus Rodriguez-Manzano, Pantelis Georgiou

**Affiliations:** Centre for Bio-Inspired Technology, Institute of Biomedical Engineering, Department of Electrical and Electronic Engineering, Imperial College London, London, SW7 2AZ, UK

## Abstract

Research into improving methods for absolute quantification of nucleic acids using standard curves has plateaued despite its positive, far-reaching impact on biomedical applications and clinical diagnostics. Currently, the mathematics involved in this mature area is restricted by the simplicity of conventional standard curves such as the gold standard cycle-threshold (*C_t_*) method. Here, we propose a novel framework that expands current methods into multidimensional space and opens the door for more complex mathematical techniques, signal processing and machine learning to be implemented. The heart of this work revolves around two new concepts: the multidimensional standard curve and its home - the feature space. This work has been validated using phage lambda DNA and standard qPCR instruments. We show that the capabilities of standard curves can be extended in order to simultaneously: enhance absolute quantification, detect outliers and provide insights into the intersection between molecular biology and amplification data. This work and its vision aims to maximise the information extracted from amplification data using current instruments without increasing the cost or complexity of existing diagnostic settings.

## INTRODUCTION

Since its inception, the real-time polymerase chain reaction (qPCR) has become a routine technique in molecular biology for detecting and quantifying nucleic acids (1-3). This is predominantly due to its large dynamic range (7-8 magnitudes), desirable sensitivity (5-10 molecules per reaction) and reproducible quantification results (4-6). New methods to improve the analysis of qPCR data are invaluable to a number of analytical fields, including environmental monitoring and clinical diagnostics (7-10).

The current “gold standard” for absolute quantification of a specific target sequence is the cycle-threshold (*C_t_*) method (11-13). The *C_t_* value is a feature of the amplification curve defined as the number of cycles in the exponential region where there is a detectable increase in fluorescence. Since this method has been proposed, several alternative methods have been developed in a hope to improve absolute quantification in terms of accuracy, precision and robustness. The focus of current research is based on the computation of single features, such as *C_y_* and –*log*_10_(*F*_0_), that are linearly related to initial concentration (14,15). This provides a simple approach for absolute quantification, however, the degrees of freedom to explore more complex data analysis techniques using multiple features are limited. Thus, research in this area has plateaued and its improvements are very incremental.

Inspired by the field of Machine Learning, this paper takes a multidimensional view, combining multiple features in order to take advantage of the information and principles behind all of the current quantification methods developed. This work describes a general framework which, for the first time, presents the multidimensional standard curve (MSC), increasing the degrees of freedom in data analysis and capable of uncovering trends and patterns in qPCR data. In fact, the conventional approach is only a special case of the proposed framework. The presented work provides a new methodology for all qPCR users in order to guarantee better quantification in any sense that the user wants, e.g. accuracy and precision.

The structure of the paper is as follows. First, the conventional approach and the proposed multidimensional framework are presented and compared. For clarity, the theory and benefits of the framework are explained and discussed in the materials and methods section. The methodology has been validated by exploring an arbitrary instance of this new framework using phage lambda DNA as a model. Here, a MSC is constructed using *C_t_, C_y_* and –*log*_10_(*F*_0_) and is shown to enhance quantification in the combination of accuracy, precision and overall predictive power. Subsequently, three qPCR assays specific to *bla*_OXA-48_, *bla*_NDM_ and *bla*_KPC_ genes are used to show the capabilities of multidimensional standard curves for outlier detection. Furthermore, temperature and primer mix concentration for the phage lambda DNA assay were altered in order to investigate patterns in the data and the robustness of the MSC for absolute quantification. Finally, a discussion section elaborates on the capabilities of this novel framework and the insights it uncovers.

**Figure 1.**
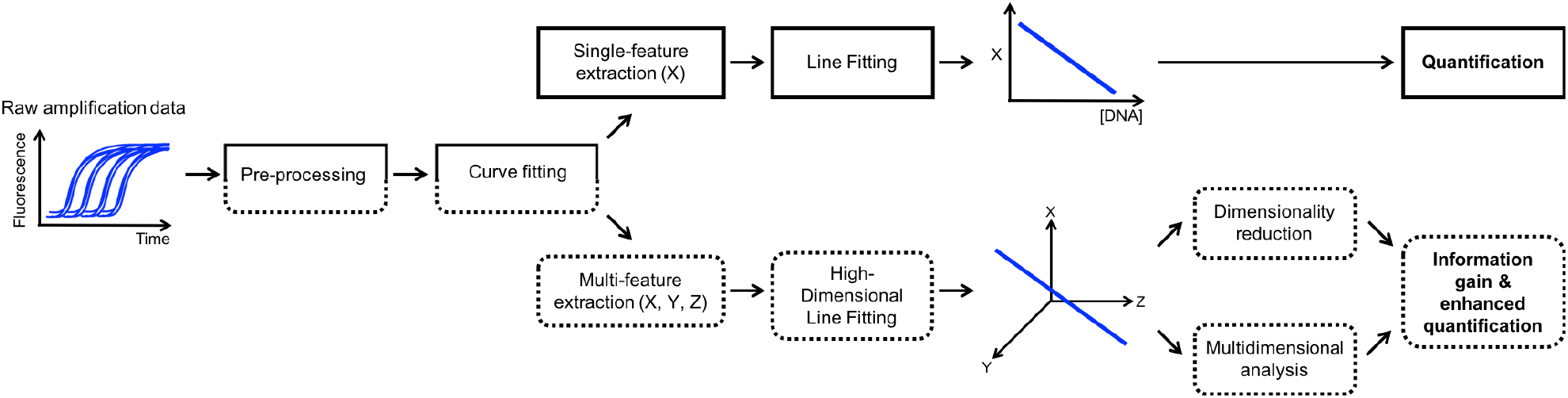
Block diagram showing the conventional method (top branch and solid line) compared to the multidimensional framework (bottom branch and dotted line) for absolute quantification. In both cases, raw amplification data for several known concentrations of the target are typically pre-processed and fitted with an appropriate curve. In the conventional case, a single feature such as the cycle threshold, *C_t_*, is extracted from each curve. A line is fit to the feature vs concentration such that unknown sample concentrations can be extrapolated. In the proposed framework, multiple features are extracted and thus a high-dimensional line is fitted in order to construct a multidimensional standard curve. Through dimensionality reduction, enhanced quantification can be achieved and using multidimensional analyses, new insights about the data can be observed.

## MATERIALS AND METHODS

### Conventional Approach

In order to understand the proposed framework, it is useful to have an overall picture of what is done in the conventional approach in the same language. Here, two terms, namely *training* and *testing* are borrowed from Machine Learning to describe the construction of a standard curve and quantifying unknown samples respectively. Within the conventional approach for quantification, training is achieved through 4 stages: pre-processing, curve fitting, linear feature extraction and line fitting. This is illustrated in Figure 1 (top branch and solid line). Testing is accomplished by using the same first 3 blocks as training, and using the line generated from the final training block in order to quantify. Pre-processing is typically necessary to tackle challenges such as background noise such that an accurate comparison amongst samples is achieved. Curve fitting is required given that amplification curves are discrete in time and most techniques require fluorescence readings that are not explicitly measured at a given time instance.

### Proposed Framework

The proposed framework extends the conventional method by increasing the dimensionality of the standard curve in order to explore, research and take advantage of using multiple features together. This new framework is presented in Figure 1 (bottom branch and dotted line). For training, there are 6 stages: pre-processing, curve fitting, multi-feature extraction, high dimensional line fitting, multidimensional analysis and dimensionality reduction. Testing follows a similar process: pre-processing, curve-fitting, multi-feature extraction, multidimensional analysis and dimensionality reduction.

Figure 2 (a-c) illustrates the idea of training and Figure 2 (d-f) shows testing using the multidimensional approach. Starting with training, Figure 2 (a) shows processed and curve-fitted real-time nucleic acid amplification curves obtained by serially diluting the known target template. In contrast with the conventional training, instead of extracting a single linear feature, multiple features denoted using the dummy labels X, Y and Z are extracted from the processed amplification curves.

Therefore, each amplification curve has been reduced to 3 values (e.g. *X*_1_, *Y*_1_ and *Z*_1_) and, consequently, can be viewed as a point in 3 dimensional space as shown in Figure 2 (b). *It is important to stress that this is a 3-D example in order to visualise the process and any number of features could have been chosen*.

**Figure 2.**
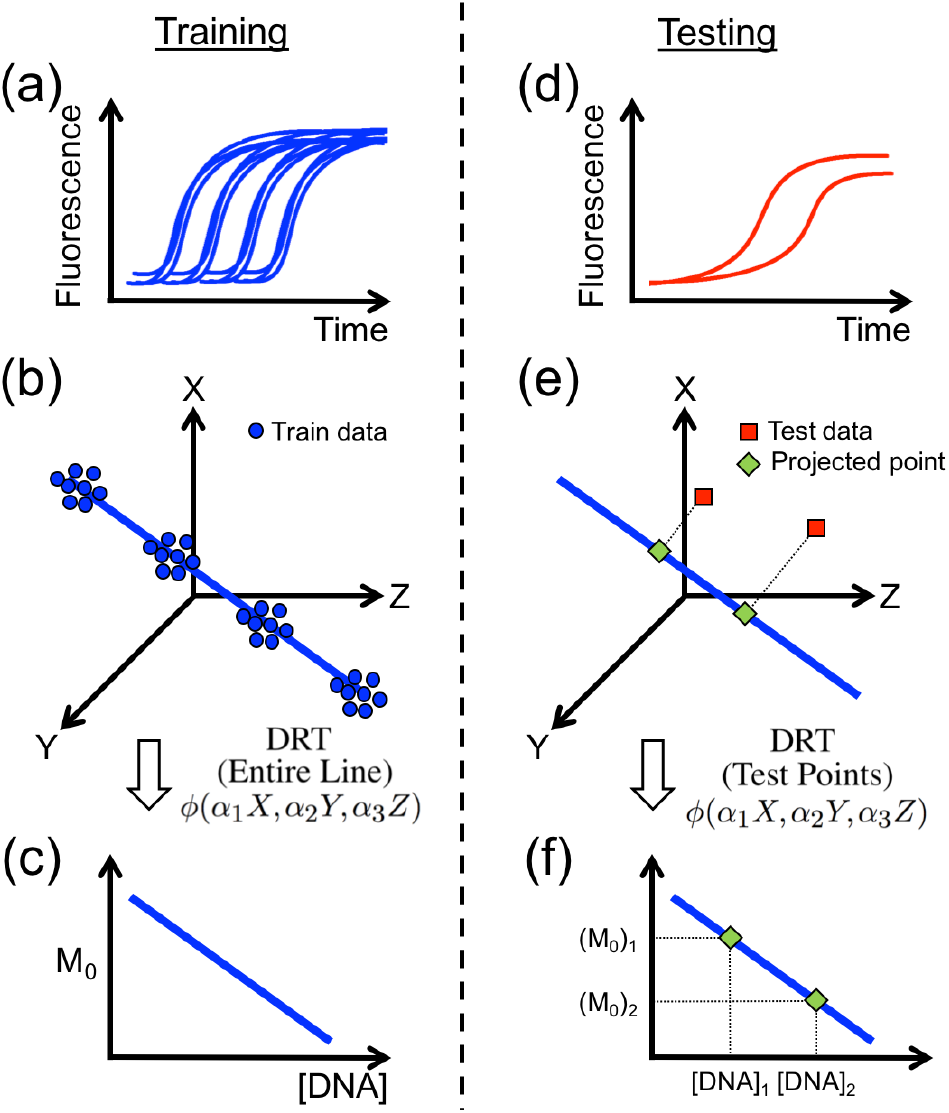
An illustration of training and testing using the multidimensional framework. (a-c) Training: (a) Processed and curve-fitted real-time amplification curves obtained from a conventional qPCR instrument using a known nucleic acid target at known concentrations. (b) Multiple features - X, Y and Z - are extracted from the processed amplification curves and plotted against each other. A multidimensional standard curve is generated through high-dimensional line fitting. (c) For quantification purposes, the MSC needs to be mapped into a quantification curve using DRTs. (d-f) Testing: (d) Unknown test amplification data is pre-processed and curve fitted. (e) The test data is projected onto the MSC. (f) Using the DRT in training, the projected data can be quantified.

In this work, only *linear* features are considered. This can be generalised to non-linear features however this is outside the scope of the proposed framework. Given that all the features are chosen such that they are linearly related to initial concentration, the training data should theoretically form a 1-D line in 3-D space. This line is approximated using high-dimensional line fitting and generates what is called the *multidimensional standard curve*. Although, the data forms a line, it is important to understand that data points do not lie exactly on the line. Consequently, there is considerable room for exploring this multidimensional space, referred to as the *feature space*, which will be reported in this paper.

For quantification purposes, the multidimensional standard curve needs to be mapped into a single dimension, defined as *M*_0_, linearly related to the initial concentration of the target. In order to distinguish this curve from conventional standard curves, it is referred to here as the *quantification curve*. This can be achieved using dimensionality reduction techniques (DRT) (16) as illustrated in Figure 2 (c). Mathematically, this means that DRTs are multivariate functions of the form: *M*_0_ = *ϕ*(*X,Y,Z*) where 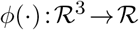. In fact, given that scaling features does not affect linearity, *M*_0_ can be mathematically expressed as *M*_0_ = *ϕ*(*α*_1_*X,α*_2_*Y,α*_3_*Z*) where *α_i_*, *i* ∈ {1,2,3} are scalar constants.

Once training is complete, unknown samples are analysed (e.g. quantified) through testing as follows. Similar to training, amplification data is processed (Figure 2 (d)) and can be considered as points in the feature space (Figure 2 (e)). Given test points may lie anywhere in the feature space, it is necessary to project them onto the multidimensional standard curve generated in training. Using the DRT function, *ϕ*, which was produced in training, *M*_0_ values for each test sample can be obtained. Subsequently, absolute quantification is achieved by extrapolating the initial concentration based on the quantification curve in Figure 2 (f).

Given that this higher dimensional space has not been reported in the literature, it is effective to highlight the degrees of freedom within this new framework that were nonexistent when observing the quantification process through the conventional lens. The following advantages arise:

*Advantage 1*. The weight of each extracted feature can be controlled by the scalars, *α*_1_,...*α_n_*. There are two main observations of this degree of freedom. The first observation is that features that have poor quantification performance can be suppressed by setting the associated *α* to a small value. This introduces a very useful property of the framework which is referred to as the *separation principle*. The separation principle means that including features to enhance multidimensional analyses does not have a negative impact on quantification performance if the *α*’s are chosen appropriately. Optimisation algorithms can be used to set the *α*’s based on an objective function (17). Therefore, the performance of the quantification using the proposed framework is lower bounded by the performance of the best single feature for a given objective. The second observation is that no upper bound exists on the performance of using several scaled features. Thus, there is a potential to outperform single features as shown in this report.

*Advantage 2*. The versatility of this multidimensional way of thinking means that there are multiple methods for dimensionality reduction such as: principal component regression, partial-least squares regression and even projecting onto a single feature (i.e. using the standard curve used in conventional methods) (18-20). Given that DRTs can be nonlinear and take advantage of multiple features, predictive performance may be improved.

*Advantage 3*. Training and testing data points do not lie perfectly on a straight line as they did in the conventional case. In fact, this property is the backbone behind why there is more information in higher dimensions. The closer two points are in the feature space, the more likely that their amplification curves are similar (resembling a Reproducing Kernel Hilbert Spaces (21)). Therefore, a distance measure in the feature space can provide a means of computing a similarity measure between amplification curves. It is important to understand that the distance measure is not necessarily, and in reality unlikely, to be linearly related to the similarity measure. For example, it is not true that a point twice as far from the multidimensional standard curve is twice as unlikely to occur. This relationship can be approximated using the training data itself. In the case of training, a similarity measure is useful to identify and remove outliers that may skew quantification performance. As for testing, the similarity measure can give a probability that the unknown data is an outlier of the standard curve, i.e. non-specific or due a qPCR artifact, without the need of post-PCR analyses such as melting curves or agarose gels (22).

*Advantage 4*. The effect of changes in reaction conditions, such as annealing temperature or primer mix concentration, can be captured by patterns in the feature space. Uncovering these trends and patterns can be very insightful in understanding the data. This is also possible in the conventional case, e.g. how *C_t_* varies with temperature, however since reaction conditions affect different features differently, conclusions can be drawn with higher confidence if a pattern is observed in multidimensional space. For example, consider the following. A change in temperature, Δ*T*, causes a different change for different features, e.g. Δ*X*, Δ*Y* and Δ*Z*. Therefore, if only a single feature, X, is used and a variation Δ*X* is observed then it is unlikely to capture the source of the variation, i.e. Δ*T* with high confidence. Whereas, considering multiple features and observing Δ*X*, Δ*Y* and Δ*Z* simultaneously provides more confidence that the source is due to Δ*T*.

An extension of advantage 4 is related to the effect of variations in target concentration. Clearly, the pattern for varying target concentration is known: along the axis of the multidimensional standard curve. Therefore, the data itself is sufficient to suggest if a particular sample is at a different concentration than another. This is significant as variations amongst replicates, which are possible due to experimental errors such as dilution and mixing, can be identified and potentially compensated for. This is of particular importance for low concentrations as the error is more significant.

Given the nature of the framework, it is now trivial to observe that the conventional approach is a special instance of the proposed framework whereby only a single feature is used. It is also interesting to observe that even if multiple features are used, if the DRT is chosen such that the multidimensional curve is projected onto a single feature, e.g. *C_t_*, then the quantification performance is exactly the same as the conventional process yet the opportunities and insights obtained in multidimensional space still remain.

### Instance of Framework

Each stage of the proposed framework can be accomplished using several different techniques. It is not the focus of this paper to explore different techniques in each stage as this is application dependent. Thus, for each stage, arbitrary methods are chosen to prove the power and versatility of this framework.

#### Pre-processing

The only pre-processing performed in this instance of framework is background subtraction. This is accomplished using baseline subtraction: removing the mean of the first 5 fluorescence readings from every amplification curve. More advanced methods can be used to improve performance (23).

#### Curve fitting

The chosen model for curve fitting is the 5-parameter sigmoid (Richards Curve) given by:

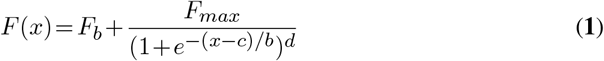

Where x is the cycle number, F(x) is the fluorescence at cycle x, *F_b_* is the background fluorescence, *F_max_* is the maximum fluorescence, c is the fractional cycle of the inflection point, b is related to the slope of the curve and d allows for an asymmetric shape (Richard’s coefficient).

The optimisation algorithm used to fit the curve to the data is the trust-region method and is based on the interior-reflective Newton method (24,25). Here, the trust-region method is chosen over the Levenberg-Marquardt algorithm (26,27) since bounds for the 5 parameters can be chosen in order to encourage a unique and realistic solution. The lower and upper bounds for the 5 parameters, [*F_b_*, *F_max_*, c, b, d], are given as: [−0.5, −0.5, 0, 0, 0.7] and [0.5, 0.5, 50, 100, 10] respectively.

#### Feature extraction

The number of features, n, that can be extracted is arbitrary, therefore 3 features were chosen in this study in order to visualise each step of the framework: *C_t_, C_y_* and –*log*_10_ (*F*_0_). Therefore each point in the feature space is a vector in 3-dimensional space, i.e. **p** =[*C_t_,C_y_*,–*log*_10_(*F*_0_)]^*T*^ where [·]^*T*^ denotes the transpose operator. Note that by convention, for the formulas in this paper, vectors are denoted using bold lowercase letters and matrices are indicated using bold uppercase letters. The details of these features are not the focus of this study thus the reader may wish to review these papers to understand each feature (14,15).

#### Line fitting

When constructing a multidimensional standard curve, a line must be fitted in n-dimensional space. This can be achieved in multiple ways such as using the first principal component in principal component analysis (PCA) or techniques robust to outliers such as random sample consensus (RANSAC (28)) if there is sufficient data. This study uses the former since a relatively small number of training points are used to construct the standard curve.

#### Distance and Similarity measure

There are two distance measures used in this study: Euclidean and Mahalanobis distance. The Euclidean distance between a point, p, and the multidimensional standard curve can be calculated by orthogonally projecting the point onto the multidimensional standard curve and then using simple geometry to calculate the Euclidean distance, e:

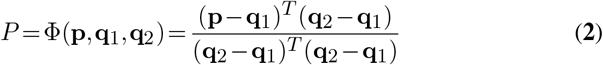

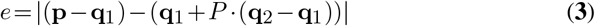

Where Φ computes the projection of the point 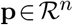 onto the multidimensional standard curve. The points **q**_1_, 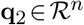 are any two distinct points that lie on the standard curve and | · | denotes the absolute value operator.

The Mahalanobis distance is defined as the distance between a point, **p**, and a distribution, 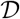, in multidimensional space (29,30). Similar to Euclidean distance, the point is first projected onto the multidimensional standard curve and the following formula is applied to compute the Mahalanobis distance, d:

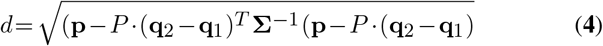

Where **p, P, q**_1_ and q_2_ are given in equation (2) and Σ is the co-variance matrix of the training data used to approximate the distribution 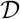.

In order to convert the distance measure into a similarity measure, it can be shown that if the data is approximately normally distributed then the Mahalanobis distance squared, i.e. *d*^2^, follows a *χ*^2^-distribution (31). Therefore, a *χ*^2^-distribution table can be used to translate a specific p-value into a distance threshold. For instance, for a *χ*^2^-distribution with 2 degrees of freedom, a p-value of 0.001 corresponds to a Mahalanobis distance of 3.72.

#### Feature weights

As mentioned previously, different weights, *α*, can be assigned to each feature. In order to accomplish this, a simple optimisation algorithm can be implemented. Equivalently, an error measure can be minimised. This is illustrated in Figure 3. In this study, the error measure to minimise is the figure of merit described in the following subsection. The optimisation algorithm is the Nelder-Mead simplex algorithm (32,33) with weights initialised to unity, i.e. beginning with no assumption on how good features are for quantification. This is a basic algorithm and only 50 iterations are used to find the weights so that there is little computational overhead.

**Figure 3.**
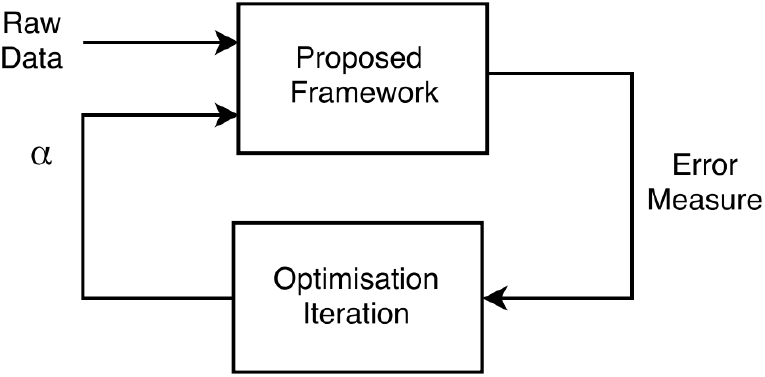
An illustration of how an optimisation algorithm can be used to find optimal parameters, *α*, for the framework.

#### Dimensionality reduction

In this study, principal component regression is used, i.e. *M*_0_ = *P* from equation (2), and it is compared with projecting the standard curve onto all three dimensions, i.e. *C_t_, C_y_* and –*log*_10_(*F*_0_).

#### Statistical Analysis

In consistency with the current literature on evaluating standard curves, relative error (RE) and coefficient of variation (CV) are used to measure accuracy and precision respectively. The CV for each concentration is calculated after normalising the standard curves such that a fair comparison across standard curves is achieved. The formula for the two measures are given by:

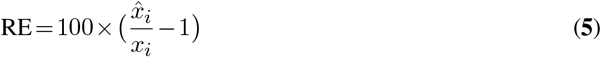

Where *i* is the index of a given training point, *x_i_* is the true concentration of the *i^th^* training data, 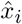 is the estimate of *x_i_* using the standard curve.

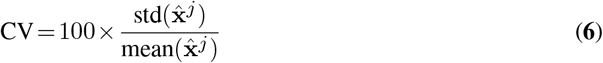

Where *j* is the index of a given concentration and 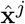 is a vector of estimated concentrations for a given concentration indexed by j. The functions *std*(·) and *mean*(·) perform the sample standard deviation and sample mean of their vector arguments respectively.

Borrowed from Statistics, this paper also uses the leave-one-out cross validation (LOOCV) error as a measure for stability and overall predictive performance (34). Stability refers to the predictive performance when training points are removed. The equation for calculating the LOOCV when a given data point is removed is given as:

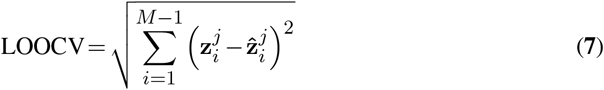

Where *M* is the total number of training data points, *i* is the index of a given training point, z_*i*_ is the true concentration of the *i^th^* training point, and 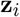 is the estimate of z_*i*_ generated by the standard curve without the *j^th^* training point. In this study, the LOOCV is specified as a percentage in order to compare across different template concentrations. This is achieved through dividing the empirical LOOCV for each concentration by its true value.

In order for the optimisation algorithm for computing *α* to simultaneously minimise the three aforementioned measures, it is convenient to introduce a figure of merit, *Q*, to capture all of the desired properties. Therefore, *Q* is defined as the product between all three errors and can be used to heuristically compare the performance across quantification methods. The average *Q* across all training data points is the error measure that the optimisation algorithm will minimise.

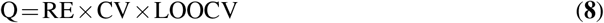

The statistical tests used to determine the significance of the results between the proposed method and conventional methods is the paired t-test for sample means with a two-sided distribution. Statistical analyses were performed in MATLAB software (The MathWorks).

### Fluorescence Datasets

Several DNA targets were used for qPCR amplification in this study:

i. Standard curves were constructed using synthetic double-stranded DNA (gblocks Fragments Genes, IDT) containing phage lambda DNA sequence (DNA concentration ranging from 10^2^ to 10^8^ copies per reaction). See supplementary data for primer and sequence information, sheet 1.
ii. Outlier detection experiments were performed using synthetic double-stranded DNA (gblocks Fragments Genes, IDT) carrying *bla*_OXA-48_, blaNDM and *bla*_KPC_ genes, in this work referred to as outlier 1, 2 and 3 respectively. See supplementary data for primer and sequence information, sheet 2.
iii. Primer and temperature variation experiments were carried out with phage lambda DNA (New England Biolabs, Catalog #N3011S) at 3 × 10^6^ copies per reaction. Final primer concentration ranged from 25 nM/each to 850 nM/each and annealing temperature ranged from 52°*C* to 72°*C*.

All oligonucleotides used in this study were synthesised by IDT (Integrated DNA Technologies, Germany) with no additional purification. The specific PCR primers for lambda phage were designed in-house using Primer3 (http://biotools.umassmed.edu/bioapps/primer3_www.cgi), whereas the primers pairs used for the outlier detection were taken from Monteiro *et al*. 2012 (35). Real-time PCR amplifications were conducted using FastStart Essential DNA Green Master (Roche) according to the manufacturer’s instructions, with variable primer concentration and a variable amount of DNA in a 5 *μL* final reaction volume. Thermocycling was performed using a LightCycler 96 (Roche) initiated by a 10 min incubation at 95°*C*, followed by 40 cycles: 95°*C* for 20 sec; 62°*C* (for lambda) or 68°*C* (for the outliers) for 45 sec; and 72°*C* for 30 sec, with a single fluorescent reading taken at the end of each cycle. Each reaction combination, starting DNA and specific PCR amplification mix, was conducted in octuplicate (5 *μL* per reaction). All the runs were completed with a melting curve analysis to confirm the specificity of amplification and lack of primer dimer. The concentrations of all DNA solutions were determined using a a Qubit 3.0 fluorometer (Life Technologies). Appropriate controls were included in each experiment.

## RESULTS

Given that there is a separation principle between quantification performance and insights in the feature space, this section is split into two parts: quantification performance and multidimensional analysis. The first part shows the results that arose from the two degrees of freedom introduced in advantage 1 & 2 and the latter explores advantage 3 & 4 regarding interesting observations in multidimensional space.

### Quantification Performance

Synthetic phage lambda dsDNA was used to construct a multidimensional standard curve and evaluate its quantification performance relative to single feature methods. The resulting multidimensional standard curve, constructed using the features *C_t_, C_y_* and –*log*_10_(*F*_0_), is visualised in Figure 4 (a). The computed features and curve-fitting parameters for each amplification curve grouped by concentration, ranging from 10^2^ to 10^8^, is presented in supplementary data, sheet 3. For comparison, Figure 4 (b) shows the quantification curves for all methods including *M*_0_ which is obtained after dimensionality reduction through principal component regression.

The optimal *α* to control the contribution of each feature to quantification, after 50 iterations of the optimisation algorithm, converged to *α* =[1.3310,0.9153,0.6386] where the weights correspond to *C_t_, C_y_* and –*log*_10_(*F*_0_) respectively. This result is readily interpretable and it suggests that –*log*_10_ (*F*_0_) exhibits the poorest quantification performance amongst the three features; as consistent with the literature (14). It is important to stress again that although the weight of –*log*_10_ (*F*_0_) is smaller than the other features to improve quantification, there is still a lot of value in keeping it as it can uncover trends in multidimensional space: as will become apparent later.

The performance measures and figure of merit, Q, for this particular instance of the proposed framework against the conventional instance is given in Table 1. A breakdown of each calculated error grouped by concentration is provided in supplementary data, sheet 4. It can be observed that in terms of the figure of merit, *M*_0_ enhances quantification by 15.8%, 29.3% and 99.6% compared to *C_t_, C_y_* and –*log*_10_ (*F*_0_) respectively. A statistical analysis was performed, as summarised in Figure 5, and the significance of the results is confirmed (p-values < 0.05).

**Table 1.** Comparison between quantification methods used in this study along with a heuristic figure of merit, Q.

| | RE (%) | CV (%) | LOOCV (%) | Fig. of Merit, $Q$ |
| --- | --- | --- | --- | --- |
| $C_t$ | $5.72 \pm 3.73$ | $0.83 \pm 0.74$ | $5.89 \pm 3.79$ | $23.37 \pm 21.01$ |
| $C_y$ | $5.97 \pm 4.08$ | $0.99 \pm 1.35$ | $6.14 \pm 4.15$ | $27.83 \pm 32.22$ |
| $F_0$ | $20.51 \pm 14.20$ | $8.65 \pm 13.76$ | $21.21 \pm 14.68$ | $5035.73 \pm 10804.56$ |
| $M_0$ | $5.77 \pm 6.06$ | $0.71 \pm 0.63$ | $5.94 \pm 3.77$ | $19.67 \pm 15.58$ |

Values are given as average ± standard deviation across concentrations with 10-fold dilutions from $10^8$ to $10^2$ . **RE** = relative error, **CV** = coefficient of variation, **LOOCV** = leave-one-out cross validation corresponding to accuracy, precision and overall predictive power respectively.

**Figure 4.**
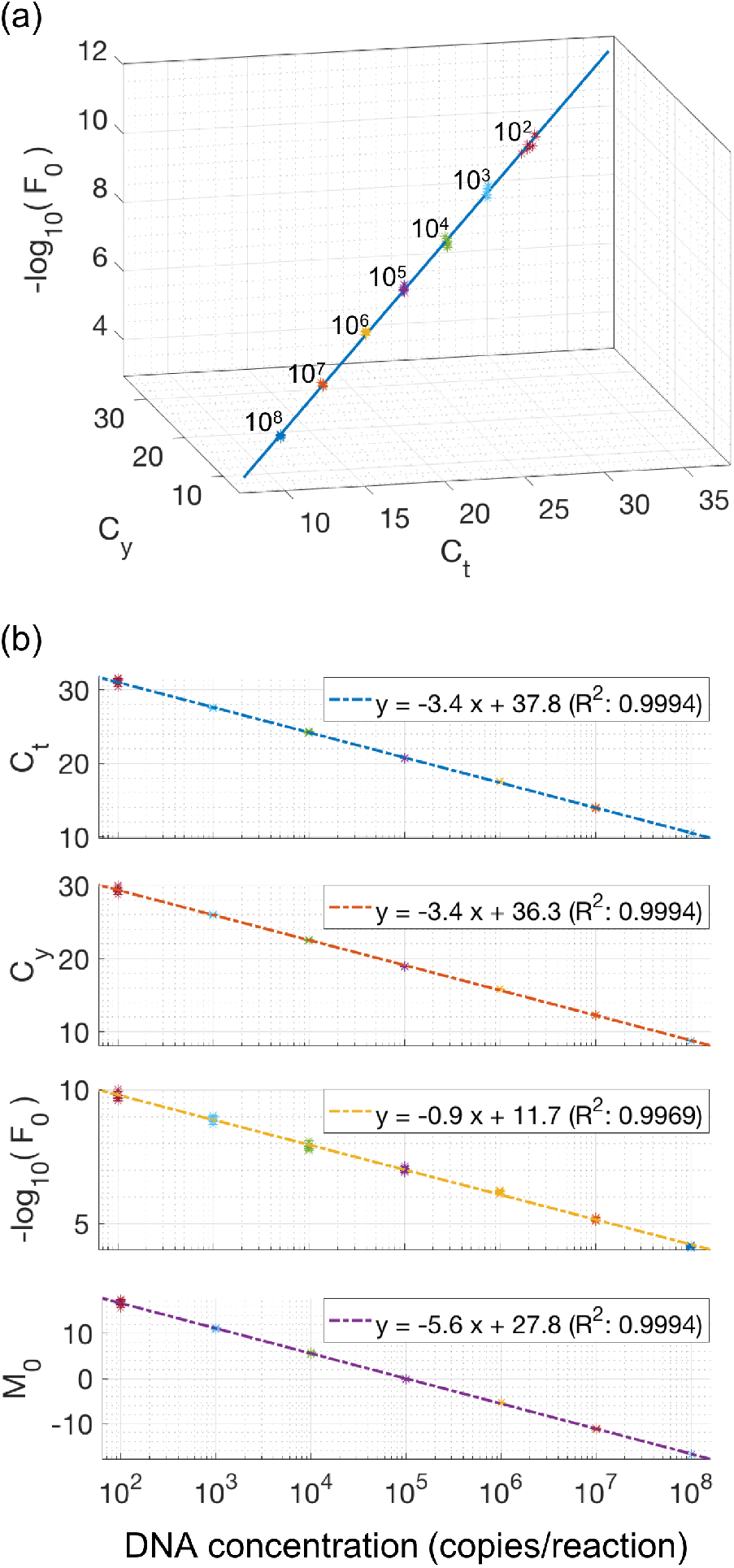
The multidimensional standard curve and quantification using information from all features. (a) A multidimensional standard curve is constructed using *C_t_, C_y_* and –*log*_10_ (*F*_0_) for lambda DNA with concentration values ranging from 10^2^ to 10^8^ (top right to bottom left). Each concentration was repeated 8 times. The line fitting was achieved using principal component analysis. (b) The quantification curve presented is obtained by dimensionality reduction of the multidimensional standard curve using principal component regression.

**Figure 5.**
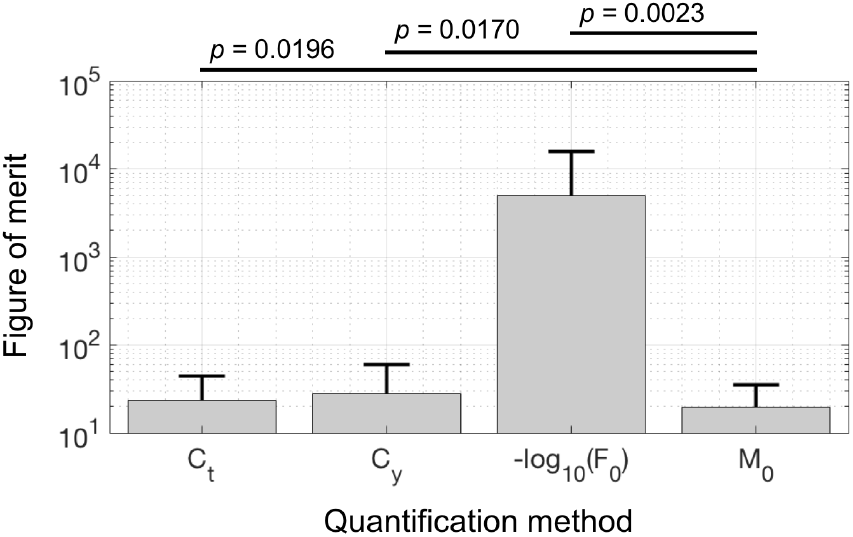
Statistical analysis for the average figure of merit combining accuracy, precision and overall predictive power. P-values between *M*_0_ and other methods are shown above the bar chart and are computed using a paired t-test with two-tail distribution.

### Multidimensional Analysis

Given the feature space is a new concept, there is a lot of room to explore what can be achieved. In this section the concept of distance in the feature space is explored and is demonstrated through an example of outlier detection. Furthermore, it is shown that a pattern exists in the feature space when altering reaction conditions.

Here, synthetic dsDNA carrying carbapenamase genes, namely *bla*_OXA-48_, *bla*_NDM_ and *bla*_KPC_, are used as deliberate outliers for the multidimensional standard curve and are referred as outlier 1, 2 and 3 respectively. Figure 6 shows the mean of the outliers in the feature space. The computed features and curve-fitting parameters for outlier amplification curves are presented in supplementary data, sheet 5. Specificity of the outliers is confirmed using a melting curve analysis as presented in supplementary data, sheet 6. Given that the outlier test points do not lie exactly on the multidimensional standard curve, Figure 6 also shows the orthogonal projection of the mean of the outliers onto the standard curve; as described in the proposed framework.

In order to fully capture the position of the outliers in the feature space, it is convenient to view the feature space along the axis of the multidimensional standard curve. This is possible by projecting data points in the feature space onto the plane perpendicular to the standard curve as illustrated in Figure 7 (a). The resulting projected points are shown in Figure 7 (b). It can be observed that all three outliers can be clustered and clearly distinguished from the training data. Furthermore, the Euclidean distance, *e*, from the multidimensional standard curve to the mean of the outliers is given by *e*_1_ = 1.16, *e*_2_ =0.77 and *e*_3_ = 1.41. Given that the furthest training point from the standard curve in terms of Euclidean distance is 0.22: the ratio between *e*_1_, *e*_2_, *e*_3_ and 0.22 is given by 5.27, 3.5, 6.41 respectively. Therefore, this ratio can be used as a similarity measure and the three clusters could be classified as outliers. However, this similarity measure has two implicit assumptions: (i) The data follows a uniform probability distribution. That is, a point twice as far is twice as likely to be an outlier. This assumption is typically made when there is not enough information to infer a distribution. (ii) Distances in different directions are equally likely. This is intuitively untrue in the feature space because a change in one direction, e.g. *C_t_*, does not impact the amplification curve as much as another, e.g. –*log*_10_(*F*_0_). It is important to emphasise that directions in the feature space contain information regarding how much amplification kinetics change and, therefore, direct comparisons between amplification reactions should be made along the same direction. This information is not captured in the conventional approach which is based on unidimensional data analysis.

**Figure 6.**
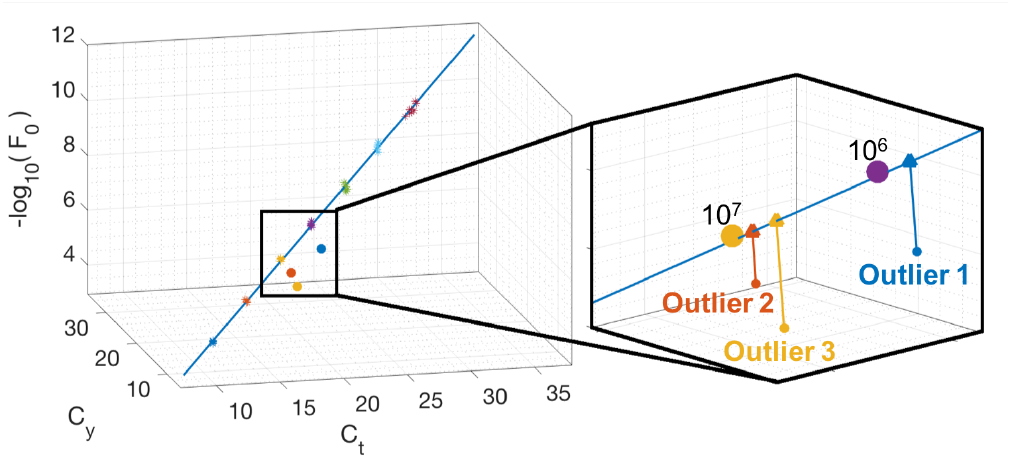
Outliers in the feature space. (left) The multidimensional standard curve for lambda DNA along with three outliers. (right) Zoomed into the region of the feature space with the mean of the replicates and the projection of the outliers onto the standard curve.

**Figure 7.**
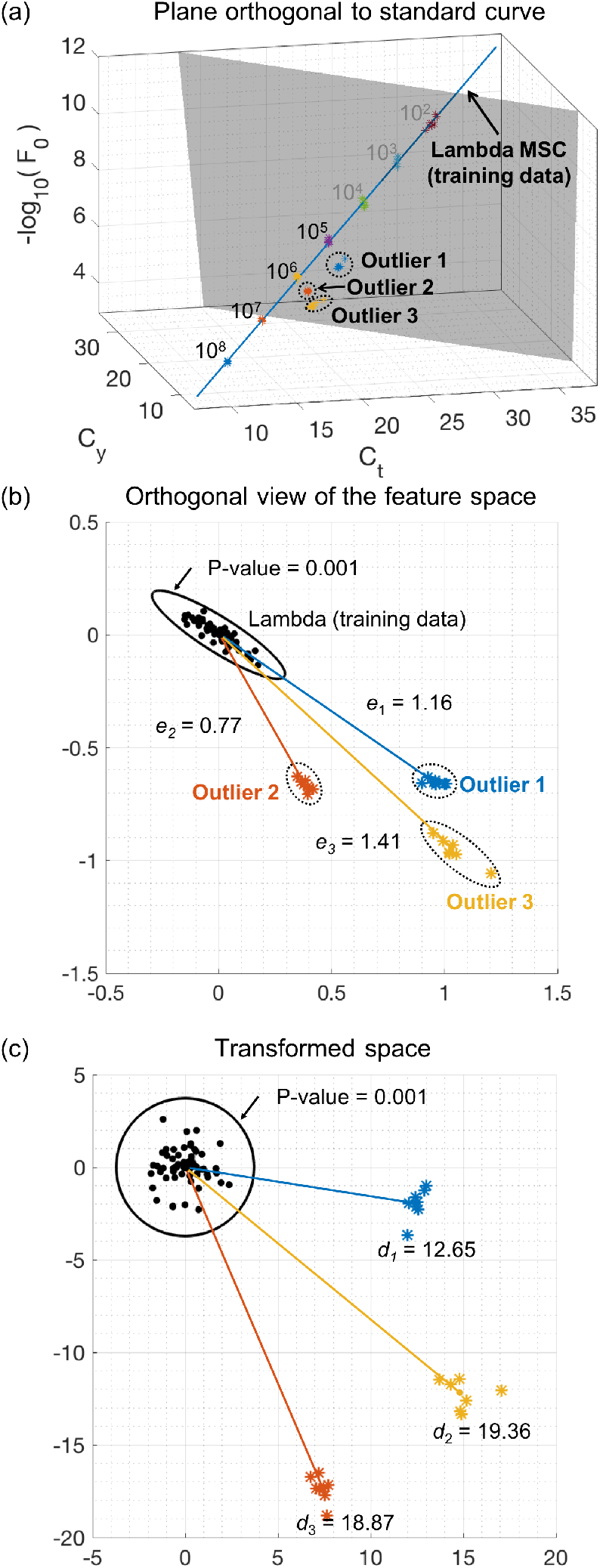
Multidimensional analysis using the feature space for clustering and detecting outliers. (a) A multidimensional standard curve using *C_t_, C_y_* and –*log*_10_ (*F*_0_) for lambda DNA with concentration values ranging from 10^2^ to 10^8^ (top right to bottom left). An arbitrary hyperplane orthogonal to the standard curve is shown in grey. (b) The view of the feature space when all the data points have been projected onto the aforementioned hyperplane. The data points consist of training standard points and the three outliers. Errors corresponding to the Euclidean distance, *e*, from the multidimensional standard curve to the mean of the outliers is given by *e*_1_ = 1.16, *e*_2_ =0.77 and *e*_3_ = 1.41. The 99.9% confidence corresponding to a p-value of 0.001 is shown with a solid black line. (c) A transformed space where the Euclidean distance, d, is equivalent to the Mahalanobis distance in the orthogonal view. The black circle corresponds to a p-value of 0.001.

In order to tackle the two aforementioned assumptions, the Mahalanobis distance, *d*, can be used. Clearly, by observing Figure 7 (b), the training data predominantly varies in a given direction. The Mahalanobis distance can be computed directly using equation (4). In order to visualise the Mahalanobis distance, the orthogonal view of the feature space (Figure 7 (b)) can be transformed into a new space (Figure 7 (c)) where the Euclidean distance is equivalent to the Mahalanobis distance in the original space. This is achieved by normalising principal components of the training data. It is now obvious from Figure 7 (c) that data in all directions are equiprobable, i.e. the training data forms a circular distribution in the transformed space. The Mahalanobis distance from the multidimensional standard curve to the mean of the outliers is given by *d*_1_ = 12.65, *d*_2_ = 18.87 and *d*_3_ = 19.36. In comparison to the Euclidean distances, it is observed that when considering the distribution of the data, the position of the outliers significantly change. As an example, based on Euclidean distance, outlier 2 is the closest whereas using the Mahalanobis distance suggests outlier 1.

A useful property of the Mahalanobis distance is that its squared value follows a *χ*^2^ -distribution if the data is approximately normally distributed. Therefore, the distance can be converted into a probability in order to capture the non-uniform distribution. Figure 8 shows the histogram of Mahalanobis distance squared for the entire training set superimposed with a *χ*^2^-distribution with 2 degrees of freedom. Based on the *χ*^2^-distribution table, any point further than 3.717 is 99.9% (p-value < 0.001) likely to be an outlier. Since all the outliers have a Mahalanobis distance significantly greater than 3.717, they are classified as outliers.

The second multidimensional analysis is concerned with observing patterns and robustness of the MSC for absolute quantification in non-ideal reaction conditions. As an example, annealing temperature and primer mix concentration are chosen to illustrate the idea. Specificity of the qPCR is not affected, as shown with melting curve analyses (see supplementary data, sheet 6). Figure 9 (a) shows the effect of how a given concentration of the phage lambda target moves in the feature space as annealing temperature is varied. It is observed that temperatures ranging from 52.0 to 69.9°*C* mostly affect –*log*_10_(*F*_0_) whereas changes from 69.9 to 72.0°*C* affects mostly *C_t_* and *C_y_* (see supplementary data, sheet 7). Similarly, Figure 9 (b) shows, that for a given concentration of phage lambda DNA, there is a pattern in the feature space associated with varying primer mix concentration from 25 to 850 nM. The pattern is observed to be approximately linear and to predominantly change along the –*log*_10_(*F*_0_) direction (see supplementary data, sheet 8). Both experiments suggest that *C_t_* and *C_y_* are more robust to changes in annealing temperature and primer mix concentration which is good for quantification performance. Furthermore, the patterns are observed in the feature space predominantly due to –*log*_10_(*F*_0_). Based on this finding, the unidimensional way of thinking would suggest to use *C_t_* or *C_y_* for future experiments. However, this implies a loss of information contained in patterns generated by –*log*_10_(*F*_0_). Therefore, the proposed multidimensional approach combines features that are beneficial for quantification performance *and* pattern recognition: preserving *all* information whilst uncompromising the quantification performance.

**Figure 8.**
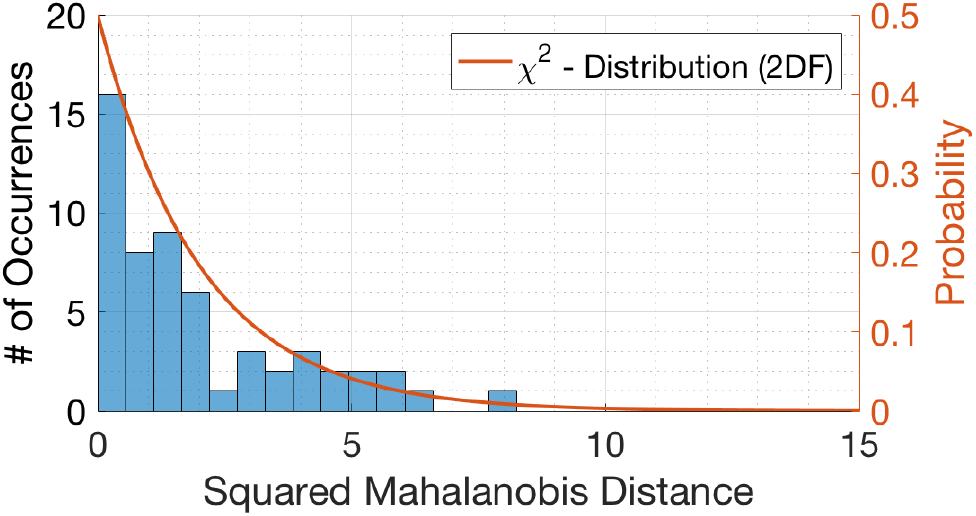
Data distribution. A histogram of the Mahalanobis distance squared of all training data points used in constructing the multidimensional standard curve superimposed with a *χ*^2^-distribution with 2 degrees of freedom (orange line).

Following this, the robustness of *M*_0_ for quantification is evaluated compared to the underlying single features. Table 2 and 3 show the estimated quantification for lambda phage DNA using each method as temperature and primer mix concentrations are varied respectively. The assumed concentration of the lambda phage DNA is 3 × 10^6^ copies/reaction; based on Qubit 3.0 fluorometer. The significance between the average quantification between methods is determined using a paired t-test with a two-tailed distribution.

For temperature variation (Table 2), the estimated average concentration for *M*_0_ is closer to the true concentration. Statistical analyses show that the results is significant compared with *C_t_* and *C_y_* (p-value < 0.1). As compared with –*log*_10_ (*F*_0_), the test is inconclusive because the variance of quantification using –*log*_10_ (*F*_0_) is very large. For primer mix concentration variation (Table 3), the estimated average concentration for *M*_0_ is comparable with *C_t_* and closer to the true concentration compared with *C_y_* and –*log*_10_ (*F*_0_). The statistical analyses show that there is no difference in the mean of *M*_0_ and *C_t_* (p-value ≈ 0.12), whereas the results are significant compared with *C_y_* and –*log*_10_ (*F*_0_) (p-value < 0.1). All data and analyses regarding variation experiments can be found in supplementary data, sheet 7 & 8. Overall, *M*_0_ outperforms or provides comparable robustness in quantification under temperature and primer mix concentration variation compared to the best single feature method.

Finally, a further interesting observation is that for low concentrations of nucleic acids, there is a variation of training data points along the axis of the multidimensional standard curve as seen in Figure 9 (c). Thus it is intuitive to hypothesise that the variation is due to fluctuations in concentration as apposed to changes in reaction kinetics. There are two implications of this assumption: (i) all the points are inliers and thus likely to be specific without the need of resource consuming post-PCR analyses. Specificity is confirmed using a melting curve analysis given in supplementary data, sheet 6. (ii) The outcome of absolute quantification is based on 3 features as apposed to a single feature which implies an increased confidence in the estimated target concentration.

**Table 2.**
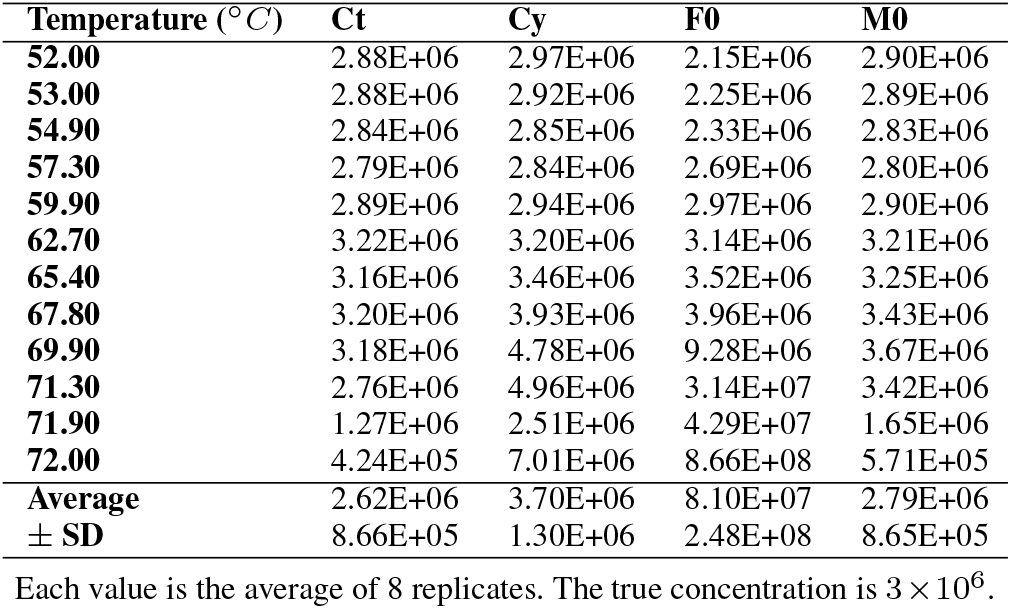
Estimated concentration when varying annealing temperature.

| Temperature ($^{\circ}\text{C}$) | $C_t$ | $C_y$ | $F_0$ | $M_0$ |
| --- | --- | --- | --- | --- |
| 52.00 | 2.88E+06 | 2.97E+06 | 2.15E+06 | 2.90E+06 |
| 53.00 | 2.88E+06 | 2.92E+06 | 2.25E+06 | 2.89E+06 |
| 54.90 | 2.84E+06 | 2.85E+06 | 2.33E+06 | 2.83E+06 |
| 57.30 | 2.79E+06 | 2.84E+06 | 2.69E+06 | 2.80E+06 |
| 59.90 | 2.89E+06 | 2.94E+06 | 2.97E+06 | 2.90E+06 |
| 62.70 | 3.22E+06 | 3.20E+06 | 3.14E+06 | 3.21E+06 |
| 65.40 | 3.16E+06 | 3.46E+06 | 3.52E+06 | 3.25E+06 |
| 67.80 | 3.20E+06 | 3.93E+06 | 3.96E+06 | 3.43E+06 |
| 69.90 | 3.18E+06 | 4.78E+06 | 9.28E+06 | 3.67E+06 |
| 71.30 | 2.76E+06 | 4.96E+06 | 3.14E+07 | 3.42E+06 |
| 71.90 | 1.27E+06 | 2.51E+06 | 4.29E+07 | 1.65E+06 |
| 72.00 | 4.24E+05 | 7.01E+06 | 8.66E+08 | 5.71E+05 |
| Average | 2.62E+06 | 3.70E+06 | 8.10E+07 | 2.79E+06 |
| ± SD | 8.66E+05 | 1.30E+06 | 2.48E+08 | 8.65E+05 |

Each value is the average of 8 replicates. The true concentration is $3 \times 10^6$ .

**Table 3.** Estimated concentration when varying primer mix concentration.

| Primer Conc. (nM) | $C_t$ | $C_y$ | $F_0$ | $M_0$ |
| --- | --- | --- | --- | --- |
| 25 | 4.45E+06 | 3.33E+06 | 3.59E+05 | 3.94E+06 |
| 100 | 3.56E+06 | 2.78E+06 | 1.19E+06 | 3.25E+06 |
| 175 | 3.25E+06 | 2.84E+06 | 3.02E+06 | 3.11E+06 |
| 250 | 2.84E+06 | 3.02E+06 | 5.66E+06 | 2.91E+06 |
| 325 | 2.43E+06 | 3.24E+06 | 2.52E+07 | 2.67E+06 |
| 400 | 2.84E+06 | 3.09E+06 | 4.15E+06 | 2.93E+06 |
| 475 | 2.73E+06 | 3.22E+06 | 5.02E+06 | 2.90E+06 |
| 550 | 2.64E+06 | 3.45E+06 | 6.94E+06 | 2.88E+06 |
| 625 | 2.72E+06 | 3.37E+06 | 5.66E+06 | 2.93E+06 |
| 700 | 2.98E+06 | 3.42E+06 | 4.02E+06 | 3.11E+06 |
| 775 | 2.79E+06 | 3.25E+06 | 4.17E+06 | 2.94E+06 |
| 850 | 2.74E+06 | 3.23E+06 | 4.42E+06 | 2.90E+06 |
| Average | 3.00E+06 | 3.19E+06 | 5.82E+06 | 3.04E+06 |
| ± SD | 5.42E+05 | 2.15E+05 | 6.37E+06 | 3.18E+05 |

Each value is the average of 8 replicates. The true concentration is $3 \times 10^6$ .

**Figure 9.**
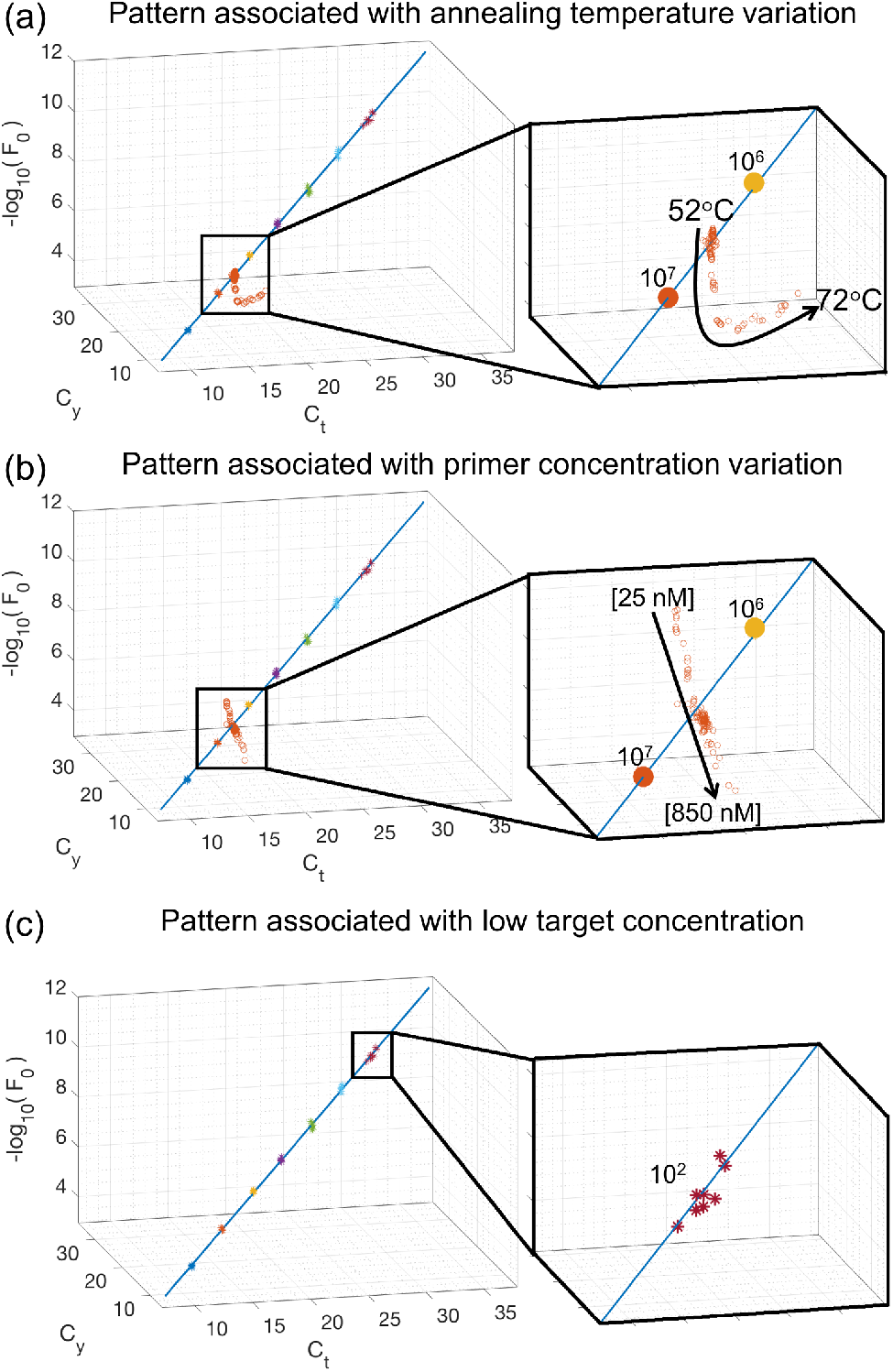
Patterns associated with changing reaction conditions. The multidimensional standard curve in all plots are using *C_t_, C_y_* and –*log*_10_ (*F*_0_) for lambda DNA with concentration values ranging from 10^2^ to 10^8^ copies/reaction (top right to bottom left). (a) The magnified image shows the effect of changing the reaction temperature from 52° *C* to 72° *C* for lambda DNA at 5 × 10^6^ copies/reaction. (b) The magnified image shows the effect of changing the primer mix concentration from 25nM to 850nM for each primer for lambda DNA at 5 × 10^6^ copies/reaction. (c) The magnified image shows the individual training sample location in the feature space for a given low concentration: 10^2^ copies/reaction.

## DISCUSSION

Absolute quantification of nucleic acids in real-time PCR using standard curves is undoubtedly important and significant in various fields of biomedicine, although research has saturated in recent years. This is partially due to the movement of research towards digital PCR (dPCR) because of the advantages it poses over qPCR such as the absence of a standard curve for absolute quantification. However, dPCR is currently not suitable for point-of-care applications given the cost and complexity of instruments. This paper presents a framework that shows the benefits of standard curves extend beyond absolute quantification when observed in a multidimensional environment. Consequently, this work opens the door for researchers from different fields to explore mathematical methods and applications that are enabled by the proposed framework.

The focus of current researchers is on the computation of a single value, referred to here as a feature, that is linearly related to concentration. Therefore, there has been a gap in the literature in taking advantage of multiple features. The potential reason for a lack of research in this area is because of the non-trivial benefits of combining linear features. The only intuitive interpretation of using several features is in the reliability of quantification. For example, instead of trusting a single feature, e.g. *C_t_*, other features such as *C_y_* and –*log*_10_ (*F*_0_) can be used to check if the quantification result is similar. This unidimensional way of thinking prevents several degrees of freedom and advantages that the proposed versatile framework enables.

There are four main capabilities that are enabled by the proposed framework: (i) the ability to select multiple features and weight them based on quantification performance. (ii) the flexibility of choosing the optimal mathematical method that maps multiple features into a single value representing target concentration. The first two capabilities lead to the separation principle which lower bounds the quantification performance of the framework to the best single feature. However, the insights and multidimensional analyses from the multiple features still remain. It is interesting to observe that for the dataset used in this study, the gold standard *C_t_* method outperformed the other single features. This is an example of why the community is reluctant to using other features given that the outcome is data dependent. The proposed framework offers a robust method of absolute quantification without the need to select a specific feature with a guaranteed quantification performance. This paper shows that in fact it is possible to increase the quantification performance as apposed to single features. (iii) the third capability enables applications such as outlier detection through the information gain captured by the elements of the feature space (e.g. distance measure, direction, distribution of data) that are typically meaningless or not considered in the unidimensional approach. (iv) the fourth advantage complements the prior advantage in that specific perturbations in reaction conditions are observed as characteristic patterns in the feature space.

The multidimensional way of thinking is not completely unfamiliar in absolute quantification. The shape based outlier detection (SOD) (36) takes a multidimensional approach in order to define a similarity measure between amplification curves. However, there are two fundamental differences with the work of this paper. The first is that SOD relies on using a specific model for amplification, namely the 5-parameter sigmoid, and is therefore not a general method. The second difference is that the pattern between the features in SOD and initial target concentration is unknown, therefore the SOD cannot be naturally integrated into the quantification process and is typically used as an add-on (37). In other words, the multidimensional approach is only considered for outlier detection and quantification is still considered as unidimensional.

The contribution of this work can be accredited to the framework as a whole and the feature space which incorporates the multidimensional standard curve. Currently, the framework is limited to considering features that are linearly related to initial target concentration. This limitation is in fact a design choice given there is a lack of other types of features available in the literature that are nonlinear and in order to reduce the complexity of the analysis. The second limitation is related to the feature space. The question arises as to whether sufficient information is captured between amplification curves in order to distinguish them in the feature space. For example, if two unrelated PCR reactions exhibit a perfectly symmetric sigmoidal amplification curve, their respective standard curves may potentially overlap. This limitation can be tackled from a molecular perspective by tuning the chemistry in order to sufficiently change amplification curves without compromising the performance of the reaction (e.g. speed, sensitivity, specificity, etc).

In terms of future directions, there are many research paths that can be explored. Both the theory of the framework and applications of the framework can be investigated. The results presented in this paper raise a number of questions: can compensating for reaction conditions significantly improve quantification performance? Does amplification efficiency describe a pattern in the feature space? Can the proposed framework be extended to non-linear features? Can the proposed framework be used for emerging isothermal amplification chemistries?

In conclusion, this paper presents a versatile framework, multidimensional standard curve and the feature space - which opens the door for researchers to explore techniques and ideas that were not previously possible. We hope by sharing these concepts, others will be able to adapt and enhance this work to meet their objectives and advance the field of nucleic acid research.

